# Instrumental aversion coding in the basolateral amygdala and its reversion by a benzodiazepine

**DOI:** 10.1101/2021.07.08.451726

**Authors:** Philip Jean-Richard-dit-Bressel, Jenny Tran, Angelos Didachos, Gavan P. McNally

**Author notes:** Correspondence to: Gavan P. McNally PhD, School of Psychology, UNSW Sydney, NSW, Australia.

## Abstract

Punishment involves learning the relationship between actions and their adverse consequences. Both the acquisition and expression of punishment learning depend on the basolateral amygdala (BLA), but how BLA supports punishment remains poorly understood. To address this, we measured calcium (Ca^2+^) transients in BLA principal neurons during punishment. Male rats were trained to press two individually presented levers for food; when one of these levers also yielded aversive footshock, responding on this punished lever decreased relative to the other, unpunished lever. In rats with the Ca^2+^indicator GCaMP6f targeted to BLA principal neurons, we observed excitatory activity transients to the footshock punisher and inhibitory transients to lever-presses. Critically, as rats learned punishment, activity around the punished response transformed from inhibitory to excitatory and similarity analyses showed that these punished lever-press transients resembled BLA transients to the punisher itself. Systemically administered benzodiazepine (midazolam) selectively alleviated punishment. Moreover, the degree to which midazolam alleviated punishment was associated with how much punished response-related BLA transients reverted to their pre-punishment state. Together, these findings show that punishment learning is supported by aversion-coding of instrumental responses in the BLA and that the anti-punishment effects of benzodiazepines are associated with a reversion of this aversion coding.

---

Our values and behaviors are shaped by our experiences. Organisms, from invertebrates to mammals, possess two highly-conserved learning systems to dynamically adapt behavior to aversive conditions: Pavlovian fear learning and instrumental aversive learning. Pavlovian fear learning allows organisms to learn predictive relationships between environmental stimuli and aversive outcomes (Stimulus–Outcome [S-O] or conditioned stimulus – unconditioned stimulus [CS-US] associations), endowing those stimuli with the ability to elicit involuntary and highly stereotyped defensive reflexes such as freezing (Blanchard & Blanchard, 1971; Rescorla, 1973). By contrast, when behavior causes aversive outcomes (punishment), we can learn the instrumental association between actions and their aversive consequences (Action-Outcome associations) to voluntarily and flexibly withhold these actions to avoid detriment (Jean-Richard-Dit-Bressel et al., 2018). Although behavioral and neural manipulation studies show that these two forms of aversive learning are distinct (Goodall & Mackintosh, 1987; Jean-Richard-Dit-Bressel et al., 2018; Jean-Richard-Dit-Bressel et al., 2019; Killcross et al., 1997b), both depend on the basolateral amygdala (BLA).

BLA has long been ascribed a critical role in Pavlovian fear learning (Maren & Quirk, 2004). Single unit recording and calcium imaging studies show that BLA principal neurons are activated by response-independent aversive outcomes (USs) (Grewe et al., 2017; Sengupta et al., 2018). This activation instructs synaptic plasticity as a substrate for Pavlovian CS-US association formation and CS salience augmentation (Sengupta et al., 2018; Wolff et al., 2014). Notably, across Pavlovian conditioning, there is a remapping of CS-evoked activity in ensembles of BLA principal neurons so that CS-evoked activity increasingly resembles US-evoked activity, with these changes in CS activity predicting the extent of CS-elicited fear reflexes (Grewe et al., 2017).

The BLA also contributes to punishment learning. Reversible inactivation or lesion of BLA impairs punishment learning and the expression of punishment avoidance independently of its role in Pavlovian conditioned fear (Jean-Richard-Dit-Bressel & McNally, 2015; Killcross et al., 1997b; Piantadosi et al., 2017). Yet how BLA activity during punishment supports punishment learning is poorly understood. This is because most existing studies of instrumental aversion have focused on active avoidance of fear CSs (Kyriazi et al., 2018) and have not isolated instrumental Action–Outcome contingencies that underpin instrumental punishment learning. So, whether punishment, like fear, is accompanied by alterations in BLA activity around punishers and their causal antecedents (i.e. punished responses) during punishment learning remains unknown.

Here we addressed these issues. We examined BLA principal neuron activity during punishment and assessed how this activity might support punishment avoidance. To do this, we used fiber photometry to measure BLA activity across a task that selectively promotes punishment learning. We also examined how anxiolytic benzodiazepines affected BLA activity during punishment. Benzodiazepines have robust, well-established anti-punishment effects via their actions as positive allosteric modulators of GABA_A_ receptors. Systemic and intra-BLA administration of benzodiazepines selectively increase punished behavior (Hodges et al., 1987; Killcross et al., 1997a; Pollard & Howard, 1990). BLA activity during punishment sessions was measured following systemic administration of benzodiazepine midazolam.

## Materials and Methods

### Subjects

Subjects were experimentally-naive male Long Evans rats (327-610g) aged 9-13 weeks at the beginning of the experiment. They were obtained from the colony maintained by UNSW Sydney, Australia. Animals were housed in groups of four in plastic cages in a climate-controlled colony room maintained on a 12hr light-dark cycle. They were given 10-15g food/day with free access to water to maintain weight at ~90% of their free feeding weight to ensure sufficient motivation to lever-press for food. All procedures were approved by the Animal Care and Ethics Committee at UNSW and conducted in accordance with the Australian Code for the Use and Care of Animals (NHMRC, 2013). Male rats were used as females tend to exhibit greater, generalized response suppression attributable to fear and not punishment learning (Jean-Richard-Dit-Bressel et al., 2019; Orsini et al., 2016), which undermines the aims of the current study.

### Apparatus

Experiments were conducted in 8 identical chambers (24 [length] × 30 [width] × 21 [height] cm; MedAssociates, St Albans, VT, USA) located inside individual light and sound attenuating cabinets (40 × 56 × 56 cm). Each chamber had stainless-steel side-walls and Perspex ceiling, back wall, and hinged door. The grid floor consisted of steel rods (4mm diameter, spaced 15mm apart) connected to a constant-current generator. The center of the right side-wall included a recess (5 x 3 x 15 cm) that housed a magazine dish (3 cm diameter) into which 45mg grain pellets (Bio-Serv, NJ, USA) were delivered. The magazine was flanked by retractable levers (2 x 4 cm). All experimental events were controlled via Med-PC IV software (Med Associates, St Albans, VT, USA). A digital camera was installed above each chamber to record animals’ spatial positions. For fiber photometry recordings, a fiber patch cable threaded into the chamber through a ceiling hole and supported by a counter-weighted gimbal holder above each chamber to minimize problematic tension and slack in the patch cable as the animal moved.

#### Optical fiber cannulae and cables

Multimode fiber optic cannulae implants and patch cables (0.39 NA, 400μm core) were manually constructed using materials from Thor Labs (Newton, NJ, USA).

#### Viral Vector

AAV vector encoding GCaMP6f driven by the CaMKIIα promoter were used (AAV5-CaMKIIα-GCaMP6f-WPRE-SV40; 1.23 x 10^13^ vp/ml) (Penn Vector Core, Philadelphia, PA, USA). The CaMKIIα promoter was used to target GCaMP6f expression to principal neurons, as CaMKIIα is predominantly expressed in BLA principal neurons and not other subtypes (McDonald et al., 2002).

#### Fiber Photometry recordings

Recordings were performed using Fiber Photometry Systems from Doric Lenses and Tucker Davis Technologies (TDT, Alachua, FL, USA). Two Doric LEDs, controlled via dual channel LED drivers, provided 465 nm (Ca^2+^-dependent signal) and 405 nm (isosbestic control signal) excitation light. GCaMP fluorescence wavelengths (~525 nm, ~ 430 nm) were measured using femtowatt photoreceivers (Newport 2151). Doric Dual Fluorescence Mini Cube (FMC2, Doric Lenses) relayed excitation/fluorescence wavelengths to/from pre-bleached patch cable and fiber optic implant. A real-time processor (RZ5P, TDT) controlled and modulated excitation lights (465 nm: 209 Hz; 405 nm: 331 Hz), as well as demodulated and low-pass filtered (3 Hz) fluorescence signals. The RZ5P also received Med-PC signals to record behavioral events in real-time. Light intensity at the tip of the patch cable was maintained at 10-40 μW across sessions. Recording fidelity was monitored in real-time; cable disconnections and other artefacts were quickly rectified and logged.

### Procedure

#### Experiment 1: Behavioral characterization of punishment

##### Lever-press Training

Rats were first given two sessions of FR1 lever-press training; both levers (left, right) were extended and pressing was reinforced with pellet delivery on an FR1 schedule (every lever-press rewarded). A lever retracted after 25 presses on it. The session ended after 1hr had elapsed or if both levers had retracted. Rats that failed to acquire lever-pressing were manually shaped in the 2^nd^ FR1 session.

Rats then received seven daily, 40min lever-press training sessions. In each session, levers were presented individually for 5min in alternating fashion; one lever was extended while the other was retracted (4 trials per lever across 40mins). Lever-pressing was reinforced with pellets on a VI30s schedule (pellet delivered on first lever-press after a variable interval [mean 30s] since last delivery). First lever to be extended (left, right) was fully randomized.

##### Punishment

Rats then received six daily, 40min punishment sessions. These sessions were identical to lever-press training, except that responses on one lever (punished lever) delivered a 0.5 sec, 0.4mA footshock on an FR10 schedule (every 10^th^ response punished). Each animal was assigned the same punished lever throughout punishment (left or right, counterbalanced across animals). The other lever remained unpunished.

##### Choice Test

Rats were then given a 10min choice test where both levers were presented concurrently. No shocks were delivered and presses on either lever delivered pellets on a shared VI60s schedule, so that there was no advantage to pressing either lever exclusively or a combination of both levers.

#### Experiment 2: BLA activity during punishment

##### Surgery

Rats were anaesthetized with 1.3 ml/kg ketamine (100 mg/ml; Ketapex; Apex Laboratories, Sydney, Australia) and 0.2 ml/kg muscle relaxant, xylazine (20 mg/ml; Rompun; Bayer, Sydney, Australia) (i.p.) and placed in stereotaxic apparatus (Model 942, Kopf, Tujunga, CA), with the incisor bar maintained at approximately 3.3 mm below horizontal to achieve a flat skull position. A 5μl, 30-gauge conical tipped microinfusion syringe (Hamilton; Reno, NV, USA) was used to inject the AAV vector (0.75μl; 0.25μl/min) into the BLA (AP −2.9, ML ±5.0, DV −8.2 mm from bregma; (Paxinos & Watson, 2007)). The syringe tip remained at the injection site for 5min to allow diffusion before being withdrawn. A 400μm optic fiber was then implanted above the BLA (AP −2.9, ML ±5.0, DV −8.0 mm from bregma) and anchored in position with dental cement (Vertex; Zeist, Netherlands) and jeweller’s screws. Immediately following surgery, animals were given i.p. injections of antibiotics (0.3ml procaine penicillin solution [300mg/ml Benicillin; Illium], 0.3ml cefazolin [100mg/ml]).

##### Behavior with fiber photometry

Behavioral training commenced 4 weeks after surgery to allow for recovery and sufficient GCaMP expression in transfected neurons. All analyzed subjects (*n* = 14) received lever-press training and 6 days of punishment, as per Experiment 1. Rats then received choice (identical to Experiment 1) and/or midazolam tests (described below); some rats received both choice and midazolam tests (*n* = 5), some only received choice test (*n* = 5), and some only received midazolam tests (*n* = 4). Thus, analyses (behavior, neural activity) for choice tests are *n* = 10, whereas analyses for midazolam tests are *n* = 9. Choice test was always conducted following punishment sessions and prior to midazolam tests, as applicable.

Subjects were connected to fiber optic patch cables and received light stimulation (405nm, 465nm) to allow BLA neural recordings for the last 3 days of lever-press training (two days for habituation, followed by one for lever-press training recording), punishment sessions, choice and midazolam tests.

##### Midazolam tests

Rats received i.p. injections of 0mg (0.9% w/v saline), 0.3mg or 1mg/ml/kg midazolam (Hypnovel, Roche, diluted with 0/9% w/v saline) fifteen minutes prior to a standard punishment session. Rats received each dose across 3 tests (within-subject, order counterbalanced). Each midazolam test day was preceded by an injection-free punishment session to limit any carry-over effects of prior sessions (including choice test).

##### Histology

At the end of the experiment, animals were anesthetized with i.p. injections of sodium pentobarbital (100mg/kg) and transcardially perfused with 0.9% saline solution containing 1% sodium nitrate and heparin (5000 IU/ml), followed by phosphate buffer solution (PB; 0.1M) with 4% paraformaldehyde. Brains were extracted, incubated in 20% sucrose solution for cryoprotection, sliced coronally (40μm) using a cryostat (Leica CM1950; Mt Waverley, Victoria, Australia), and stored in PB solution with 0.1% sodium azide at 4°C.

Fiber placement and GCaMP expression were determined using fluorescent immunohistochemistry. Brain tissue was washed in PB, incubated in PBT-X solution (10% horse serum [NHS], 0.5% Triton X-100 in PB) for 2 hours, and then incubated in PBT-X solution (2% NHS, 0.2% TritonX-100 in PB) with primary antibody (1:1000 polyclonal rabbit anti-GFP, ThermoFisher Scientific, #A11122) at room temperature for 24hr. Tissue was then washed with PB and incubated overnight in PBT-X (2% NHS, 0.2% TritonX-100 in PB) with secondary antibody (1:1000 AlexaFluor 488-conjugate anti-rabbit, ThermoFisher Scientific). Tissue was washed with PB and mounted onto gelatinised slides. Slides were left to dry and then cover-slipped using Entellan (Merck Pty Limited, Darmstadt, Germany). GCaMP6f expression and cannula placements were verified using fluorescent microscopy (Olympus BX51; Tokyo, Japan). Animals were excluded from analyses if fiber tip and GCaMP expression could not be confirmed as co-localized in BLA; *n* = 14 animals were included across analyses.

### Data Analysis

Behavior was analyzed using within-subject planned contrasts in PSY statistical software (Bird, 2004). All fiber photometry data were analyzed using custom MATLAB scripts. The Type I error rate (α) for all analyses was controlled at 0.05.

#### Behavior – lever-pressing

The primary behavioral dependent variables were lever-press rates on punished and unpunished levers. Punishment and choice test lever-press rates were analyzed using orthogonal contrasts for lever (punished vs unpunished), session (linear, quadratic) and interaction contrasts. Lever-training data was analyzed separately from punishment session data. Latencies to first lever-press (averaged across trials per session) were analyzed using the same contrasts. Lever-press rates across MDZ test were analyzed using simple effect contrasts comparing an MDZ dose (0.3mg or 1mg) against control (0mg).

In addition to raw lever-press rates, ratios of lever-pressing were used to assess self-normalized change in lever-pressing. An elevation ratio was used to compare lever-press (LP) rates under a dose of midazolam (0.3mg or 1mg MDZ) against control (0mg, i.e. saline), calculated per lever as follows:

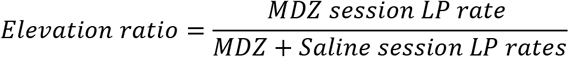

Elevation ratios are bounded from 0 to 1. A score of 0.5 indicates no change in lever-press rate following MDZ relative to saline, a score higher than 0.5 indicates increased lever-pressing relative to saline, and a score lower than 0.5 indicates decreased lever-pressing relative to saline. Elevation ratios were compared against the null ratio of 0.5 using single mean t-tests.

A suppression ratio was used to compare punished lever-press (PunLP) rates across midazolam tests (0mg, 0.3mg, 1mg) relative to pre-punishment training (T), calculated as follows:

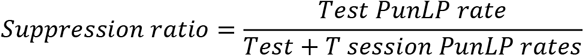

Suppression ratios are bounded from 0 to 1. A score of 0.5 indicates no difference in punished lever-press rate during an MDZ test session relative to training, a score higher than 0.5 indicates more lever-pressing relative to training, and a score lower than 0.5 indicates less lever-pressing relative to training.

#### Behavior – location and immobility

To analyze spatial position and immobility during punished and unpunished trials (periods when punished and unpunished levers were extended, respectively), video recordings from last day of lever-press training and selected punishment sessions (P1, P4, P6) were imported into EthoVision XT 10 (Noldus, Wageningen, Netherlands) to define animal center-point. To quantify spatial position, zones (4 [length] × 6 [width] cm) around the punished and unpunished levers were defined as punished and unpunished zones. The cumulative time spent in each zone was calculated in seconds. Freezing was measured as proportion of trials spent immobile. One rat was excluded from spatial location and immobility analyses due to a recording malfunction. This resulted in a final group size of *N* = 7 for analyses of spatial location and immobility, and *N* = 8 for all other analyses.

#### Fiber photometry – signal processing

Ca^2+^-dependent (465nm-related) and isobestic (405nm-related) signals and event timestamps were extracted into MATLAB, and signals during logged disconnections were discarded. Each signal was low-pass (3Hz) and band-stop filtered (1.9-2.2Hz) to remove high-frequency noise identified via Fast Fourier Transform. The isobestic signal was linearly regressed onto the Ca^2+^-dependent signal to create a fitted isobestic signal, and a normalized fluorescence change score (dF/F) was calculated using the standard formula:

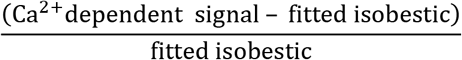

This motion-artifact-corrected dF/F was then detrended via a 600s moving average (convolution window), such that each 10min period had a mean dF/F of 0. All peri-event activity used in analyses were derived from this detrended dF/F.

#### Fiber photometry – event-related activity

The key dependent variable was BLA activity transients around aversive footshock, and punished and unpunished lever-presses. To assess these, dF/F from −3s to +7sec around lever-presses uncontaminated by outcomes (i.e. yielding footshock or pellet delivery) and footshocks (necessarily concomitant with punished lever-press) were collated. Peri-event activity kernels were obtained by normalizing each trial waveform according to its sum square deviation from 0 (Liu et al., 2020). Activity kernels were averaged per subject; all analyses used mean activity kernels per subject, i.e. was subject-based. Due to the scarcity of punished lever-presses and shock deliveries towards the end of punishment, sessions P5 and P6 were combined to obtain more accurate peri-event activity traces per subject.

To determine significant transients, 95% confidence intervals (CI) around activity kernels were derived via bootstrapping (Jean-Richard-dit-Bressel et al., 2020). Specifically, bootstrapped means were obtained by randomly resampling from subject mean waveforms with replacement (1000 iterations). CI limits were derived from 2.5 and 97.5 percentiles of bootstrap distribution, expanded by a factor of 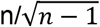 (Jean-Richard-dit-Bressel et al., 2020). A significant transient was identified as a period that CI limits did not contain 0 (moving average baseline) for at least 1/3secs (low-pass filter window; Jean-Richard-dit-Bressel et al., 2020). Punished and unpunished lever-press transients were directly compared against each other by bootstrapping the within-subject difference waveform (mean punished–mean unpunished waveform per subject).

#### Fiber photometry – kernel similarity

Similarity between lever-press and shock activity kernels across sessions were quantified by deriving a normalized fit score per kernel comparison. Fit score was calculated as the dot product of two waveform vectors (each normalized according to sum square deviation from 0). This score can range from −1 to 1. A fit above 0 indicates the two waveforms deviate from baseline in a similar way across the comparison window (1 = identical waveforms), whereas a negative fit indicates the two waveforms deviate from baseline in opposite directions (−1 = mirror opposite waveform). As each kernel is normalized, this method is specifically sensitive to differences in the specific shape of waveforms across the event window, not waveform magnitude.

To determine whether lever-press and shock waveforms across sessions were significantly similar (fit > 0) or inverse (fit < 0) to each other, 95% CI limits for fits were obtained from 2.5 and 97.5 percentiles of bootstrapped fit distributions (fits of randomly resampled mean waveforms; 1000 iterations). Two activity kernels were identified as significantly similar if the fit CI was entirely above 0, or significantly inverse if fit CI was entirely below 0.

To visualize the overall similarity/dissimilarity of activity kernels across sessions, fit scores were converted into fit distances (1 – fit [perfect fit = 0, perfect inverse = 2]). Kernel coordinates in 2D space were obtained via multidimensional scaling (MATLAB *mdscale* function, criterion = metricstress), using fit distances as input. Stress was 0.038, indicating an excellent representation of kernel similarity/dissimilarity within 2D space.

#### Fiber photometry – relative similarity of punished lever activity

To assess the relationship between BLA lever-press activity and punishment avoidance across MDZ tests, a relative fit of MDZ test lever-press activity to late punishment vs. pre-punishment was calculated per subject. Specifically, relative fit was the normalized fit (as described above) of a subject’s MDZ test punished lever-press activity against their late punishment (P5-6) punished lever-press activity, minus the normalized fit of MDZ test activity against their pre-punishment (T) punished lever-press activity (P5-6 fit – T fit). This score captures the degree a subject’s MDZ punished lever-press kernel conforms to learned punishment vs. pre-punishment kernels. A positive relative fit score indicates an MDZ test kernel is closer to late punishment than pre-punishment, whereas a negative relative fit score indicates a kernel more similar to pre-punishment.

The relationship between relative fit and suppression ratio across MDZ tests was assessed using linear regression (GraphPad Prism 9, San Diego, CA, USA).

## Results

### Experiment 1: Behavioral characterization of punishment

Rats were initially trained to respond on two individually-presented levers (5 min alternating trials) for food, each lever reinforced on a VI30s schedule. This VI30 schedule remained in effect for the remainder of the experiment. The mean ± standard error of the mean (SEM) lever-press rates at the end of training (T) are shown in Figure 2B (*left panel*). There was no difference between to-be punished and to-be unpunished lever-pressing (*F_(1,7)_* = .614, *p* = .459) (Figure 2B, *left panel* [T]). There were also no differences in latencies to initially respond on to-be punished and to-be unpunished levers (*F_(1,7)_* = .275, *p* = .616) (Figure 2B, *right panel* [T]).

Next, during punishment sessions (P1 to P6), one of these responses was punished via pairings with a 0.4 mA footshock on an FR10 schedule (Jean-Richard Dit Bressel & McNally, 2014). Animals exhibited robust punishment avoidance. They pressed the punished lever less than the unpunished lever (lever main effect: *F*_(1,7)_ = 77.079, *p* < .001). There was a lever × session interaction (linear: *F*_(1,7)_ = 34.803, *p* < .001, quadratic: *F*_(1,7)_ = 6.780, *p* = .035); punished lever-presses initially decreased followed by a modest increase across sessions (session quadratic [punished only]: *F*_(1,7)_ = 14.768, *p* = .006), whereas unpunished lever-presses increased across sessions (session linear [unpunished only]: *F*_(1,7)_ = 25.706, p <.001).

Punishment also affected average latencies to first lever-press across sessions (Figure 1B, *right panel*). Rats were slower to respond on the punished lever than the unpunished lever (*F*_(1,7)_ = 36.307, *p* = .001). A lever × session interaction was also observed (quadratic: *F*_(1,7)_ = 17.015, *p* = .004); latencies to respond on the punished lever increased and then decreased across sessions (*F*_(1,7)_ = 20.570, *p* = .003), whereas latencies to respond on the unpunished lever did not significantly change across sessions (*F*_(1,7)_ = .819, *p* = .396).

**Figure 1.**
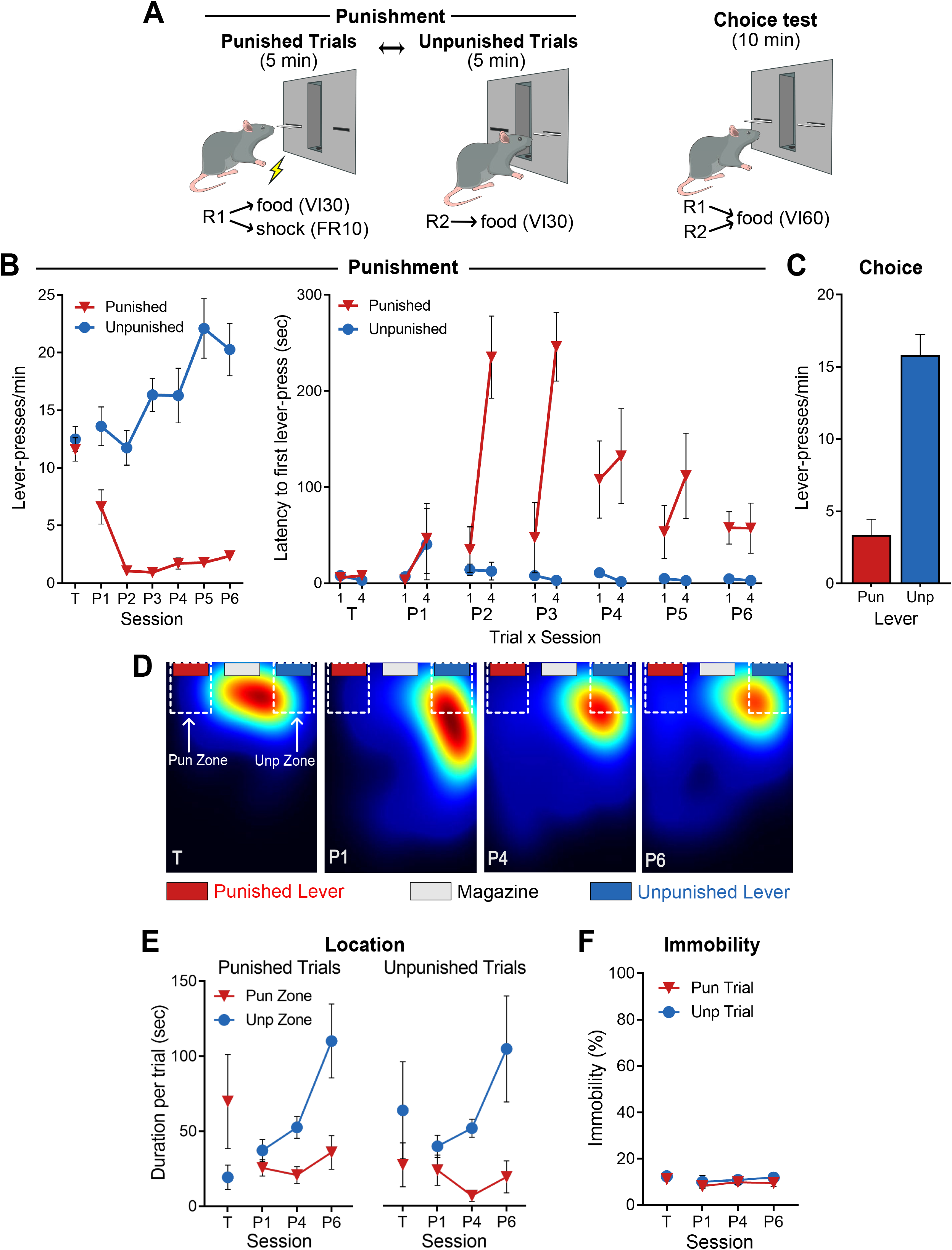
Punishment task and behavior. **[A]** Punishment task. *Left*: After training rats to press two individually-presented levers (5min alternating trials) for food (30s variable interval [VI30s] schedule), rats receive punishment sessions where presses on the punished lever also yielded footshock (FR10 schedule). *Right*: During choice test, both levers are presented to assess lever preference. **[B]** *Left*: Mean±SEM lever-press rate for last session of lever-press training (T) and across punishment sessions (P1-P6). Prior to punishment, rats (*n* = 8) responded equally on both levers. Across punishment sessions, rats suppressed punished, but not unpunished, responding. *Right*: Mean±SEM latency to first lever-press for first (1) and last (4) trial per session. Rats were slower to press the punished lever relative to unpunished lever across punished sessions, indicative of learned avoidance. **[C]** During choice test, rats preferred pressing the unpunished lever over the punished lever. **[D]** Example heatmaps of rat location during lever-press training (T) and punishment sessions (P1, P4, P6). Areas immediately around the punished and unpunished levers were designated as Pun Zone and Unp Zone, respectively. **[E]** Mean±SEM time spent in punished and unpunished zones during punished and unpunished trials (*left* and *right*, respectively) across lever-press training (T) and punishment sessions (P1, P4, P6). Before punishment, rats were preferentially located in the extended lever’s zone. During punishment, rats tended to be in the unpunished zone, regardless of which lever was extended. **[F]** Mean±SEM immobility across punished and unpunished trials. Low levels of immobility (proxy for freezing) were observed across sessions, suggesting low levels of Pavlovian fear in this task.

Finally, rats received a choice test, during which both levers were extended but neither was punished. Animals made more unpunished than punished lever-presses (*F_(1,7)_* = 70.212, *p* < .001) (Figure 1C), indicating a strong preference for unpunished over punished lever, despite the absence of shock on these tests.

We also assessed spatial location across punishment. Prior to punishment, there was no overall difference in time spent in punished versus unpunished zones (*F*_(1,6)_ = .044, *p* = .841) or time spent around levers during punished versus unpunished trials (*F*_(1,6)_ = .003, *p* = .958) (Figure 1D-E). However, there was a significant interaction between zone and trial (*F*_(1,6)_ = 6.386, *p* = .045); rats tended to stay in the zone of whichever lever was extended, although follow-up simple effect analyses on zone per trial were insignificant (all *F*_(1,6)_ ≤ 5.689, *p* ≥ .054).

Punishment changed these spatial preferences (Figure 1D-E). During punishment sessions, rats spent more time in unpunished compared to punished lever zones (*F*_(1,6)_ = 11.88, *p* = .014), and spent less time around levers during punished trials than unpunished trials (*F*_(1,6)_ = 8.875, *p* = .025). There was no trial x zone interaction (*F*_(1,6)_ = .234, *p* = .646); rats spent more time in unpunished zones during both punished (*F*_(1,6)_ = 7.208, *p* = .036) and unpunished (*F*_(1,6)_ = 7.703, *p* = .032) trials. The extent of this spatial preference for the unpunished over the punished zone increased across punishment sessions (*F*_(1,6)_ = 8.479, *p* = .027); rats increasingly spent more time around the unpunished lever (*F*_(1,6)_ = 8.753, *p* = .025), but did not significantly change time spent near the punished lever (*F*_(1,6)_ = 0.136, *p* = .725) across punishment sessions. This shows emergence of a preference for spending time near the unpunished lever, regardless of whether the punished or unpunished lever was extended. Visual observations suggested that animals spent more time exploring the chamber during punished trials, frequently returning to the unpunished zone to check for the unpunished lever.

#### Immobility

Pavlovian aversive learning instructs Stimulus-Outcome (CS-US) associations (Rescorla, 1973), which control innate behaviors such as freezing (Blanchard & Blanchard, 1971). Punishment instructs Action–Outcome associations, controlling voluntary withholding of a specific action (Bolles et al., 1980). However, involuntary, reflexive behaviors like freezing may interfere with and suppress lever-pressing, thereby confounding the measure of punishment with fear (Jean-Richard-Dit-Bressel et al., 2018). So, we directly measured immobility across punishment sessions to determine whether and when immobility may have contributed to the suppression we observed. Immobility was low across all sessions (Figure 1F). There were no differences in immobility during punished and unpunished trials prior to punishment (*F*_(1,6)_ = 1.852, *p* = .222) (Figure 1F). During punishment, there was no main effect of trial (*F*_(1,6)_ = 1.569, *p* = .257), and no trial × session interaction (*F*_(1,6)_ = .050, *p* = .830). This shows that immobility and freezing were not the cause of lever-pressing effects within this task (Jean-Richard-Dit-Bressel & McNally, 2015).

### Experiment 2: Basolateral amygdala calcium transients across punishment

Next, we next examined population-level BLA principal neuron calcium (Ca^2+^) transients during punishment learning using fiber photometry. An AAV encoding GCaMP6f under control of the CaMKIIα promoter was applied to the BLA to express the genetically-encoded Ca^2+^ sensor in BLA principal neurons (McDonald et al., 2002). GCaMP6f fluorescence was measured via an optic fiber cannula implanted in BLA (Figure 2A,C).

**Figure 2.**
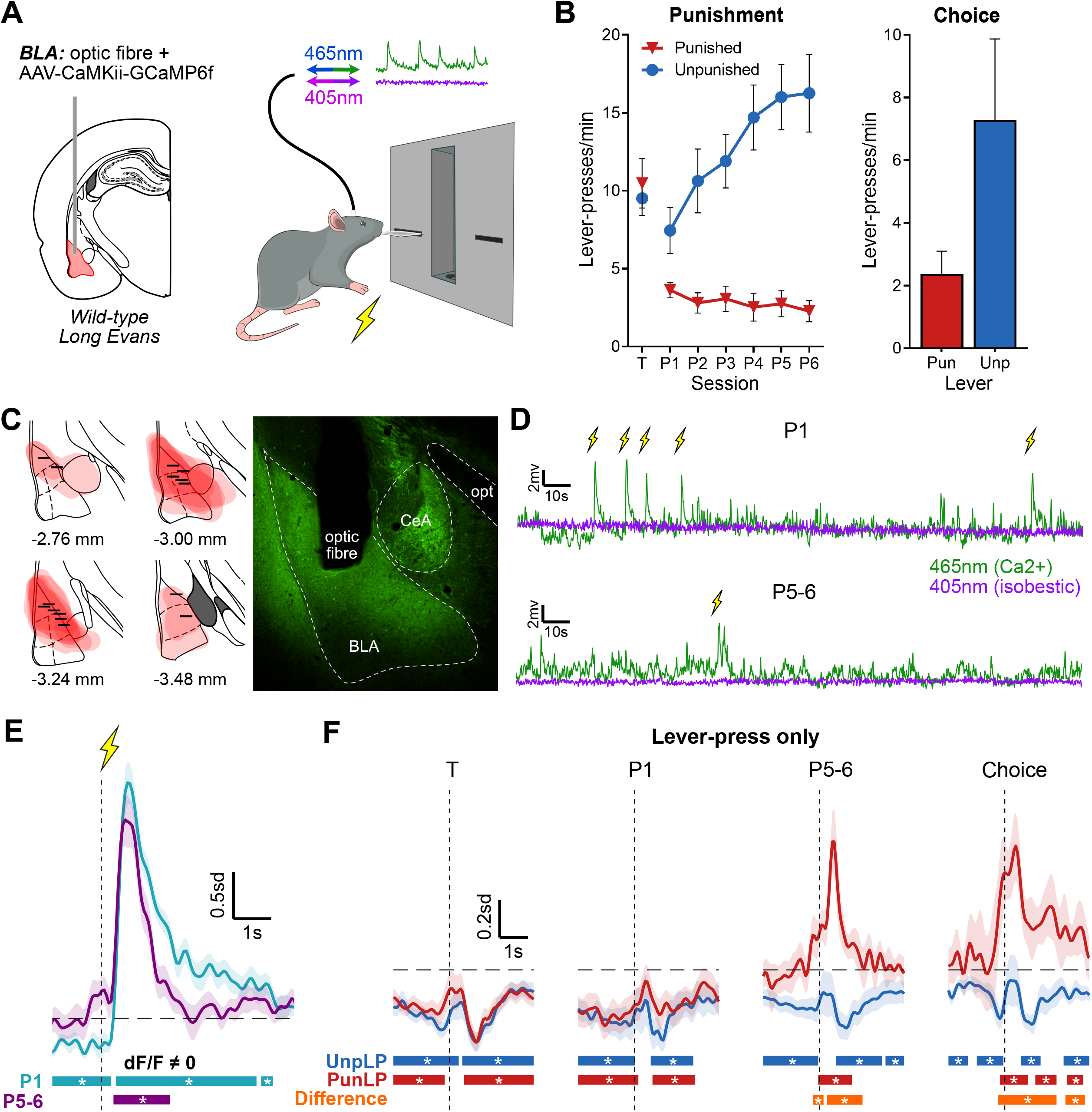
Basolateral amygdala activity during punishment. **[A]** Fiber photometry to assess basolateral amygdala (BLA) activity. *Left*: Rats received unilateral optic fiber implant and application of AAV-CaMKii-GCaMP6f into BLA. *Right*: BLA neuron activity was measured throughout punishment. **[B]** Behavior of BLA photometry animals. *Left*: Mean±SEM lever-press rate on punished and unpunished levers for last day of lever-press training (T) and punishment sessions (P1-P6) (*n* = 14). *Right*: Mean±SEM lever-press rate during choice test (*n* = 10). **[C]** *Left*: GCaMP6f expression and fiber tip location for animals included in analysis (*n* = 14). *Right*: Example BLA placement. **[D]** Example of 465nm- and 405nm-related signals during early (P1; *top*) and late (P5-6; *bottom*) punishment sessions, with shock deliveries times. **[E]** Mean±SEM of subject-based (*n* = 14) BLA activity kernel around shock deliveries during early (P1; teal) and late (P5-6; purple) punishment. Bars at bottom of graph indicate significant deviations from baseline (dF/F ≠ 0), determined via bootstrapped confidence intervals (95% CI). Vertical dashed line indicates shock onset, horizontal dashed line indicates baseline (dF/F = 0). **[F]** Mean±SEM BLA activity kernel around punished (PunLP; red) and unpunished (UnpLP; blue) lever-presses (no outcome) across training (T), punishment (P1, P5-6) and choice test. Bars at bottom of graph indicate significant deviations from baseline (95% CI) for PunLP and UnpLP, and significant differences between punished and unpunished kernels (Difference; orange). Vertical dashed lines indicate time of lever-press, horizontal dashed lines indicate baseline (dF/F = 0).

The mean ± SEM lever-press rates for final lever-press training session (T) and punishment sessions (P1 – P6) are shown in Figure 2B. There were no differences in leverpress rates for to-be punished and unpunished levers at end of lever-press training (*F*(1,13) = 0.466, *p* = 0.507). During punishment, animals pressed the unpunished lever more than punished lever (lever main effect: *F*(1,13) = 30.72, *p* < .001). This difference increased across sessions (lever × session interaction: *F*(1,13) = 22.56, *p* < .001); unpunished lever-pressing increased (*F*(1,13) = 36.31, *p* < .001) whereas punished lever-presses decreased (*F*(1,13) = 6.12, *p* = .028) across punishment. For animals that underwent choice test (*n* = 10; Figure 2B [right]), there was a significant preference for unpunished over punished levers (*F(1,9)* = 18.75, *p* = .002).

The first question was how BLA activity related to delivery of punishment. We analyzed Ca^2+^ transients using 95% confidence intervals with a consecutive threshold of 0.33s (Jean-Richard-dit-Bressel et al., 2020). The BLA exhibited robust excitatory Ca^2+^ transients to response-generated footshock across punishment (Figure 2E), showing recruitment of BLA principal neurons by the punisher. This is similar to previous reports of excitatory Ca^2+^ transients to response-independent footshocks in studies of Pavlovian fear (Grewe et al., 2017; Sengupta et al., 2018).

Punishment learning involves encoding the instrumental relationship between actions and their aversive outcomes. So, the next question was whether punishment changed BLA encoding of punished responses (lever-presses). Prior to punishment, there were significant inhibitory transients (Figure 2F [T]) associated with responses on both the to-be punished and to-be unpunished levers. This same activity pattern was observed in the first session of punishment (Figure 2F [P1]). However, when punishment learning was established (Figure 2F [P5-6]), BLA transients to punished lever-presses were no longer inhibitory. Instead, there were now significant excitatory transients around punished lever-press. Importantly, this change in BLA transients was response-specific: Ca^2+^ transients to unpunished lever-presses remained unchanged and inhibitory. Within-subject comparisons of punished and unpunished lever-press transients confirmed significant differences between activity around punished and unpunished lever-presses.

Critically, the same pattern of BLA activity around lever-presses was observed during choice test when the punisher was absent (Figure 2F; *n* = 10). Punished and unpunished lever-press transients were significantly different from each other, with punished and unpunished lever-pressing characterized by significant excitatory and inhibitory transients, respectively. This preservation of excitatory transients during choice test shows that punished lever-press transients were not dependent on recent exposure to aversive outcomes *per se* or on the trial-based structure of punishment sessions.

#### Midazolam tests

The identification of punished lever-press transients which emerged across the course of punishment training is consistent with the possibility that BLA activity encodes the learned aversive value of instrumental actions. To test this possibility more directly, we examined the effects of midazolam (MDZ) on these transients and punished behavior. Benzodiazepines have well-documented anti-punishment effects, specifically increasing punished behavior (Pollard & Howard, 1990). At higher doses, benzodiazepines can also have sedative effects, reducing behavior generally. Following punishment (Figure 2), we tested the effect of systemically administered benzodiazepine, midazolam (MDZ), on behavior and BLA activity (Figure 3). Subjects (*n* = 9) received control (0mg/kg) and MDZ (0.3 or 1mg/kg) injections prior to separated punishment sessions (within-subject, order counter-balanced).

**Figure 3.**
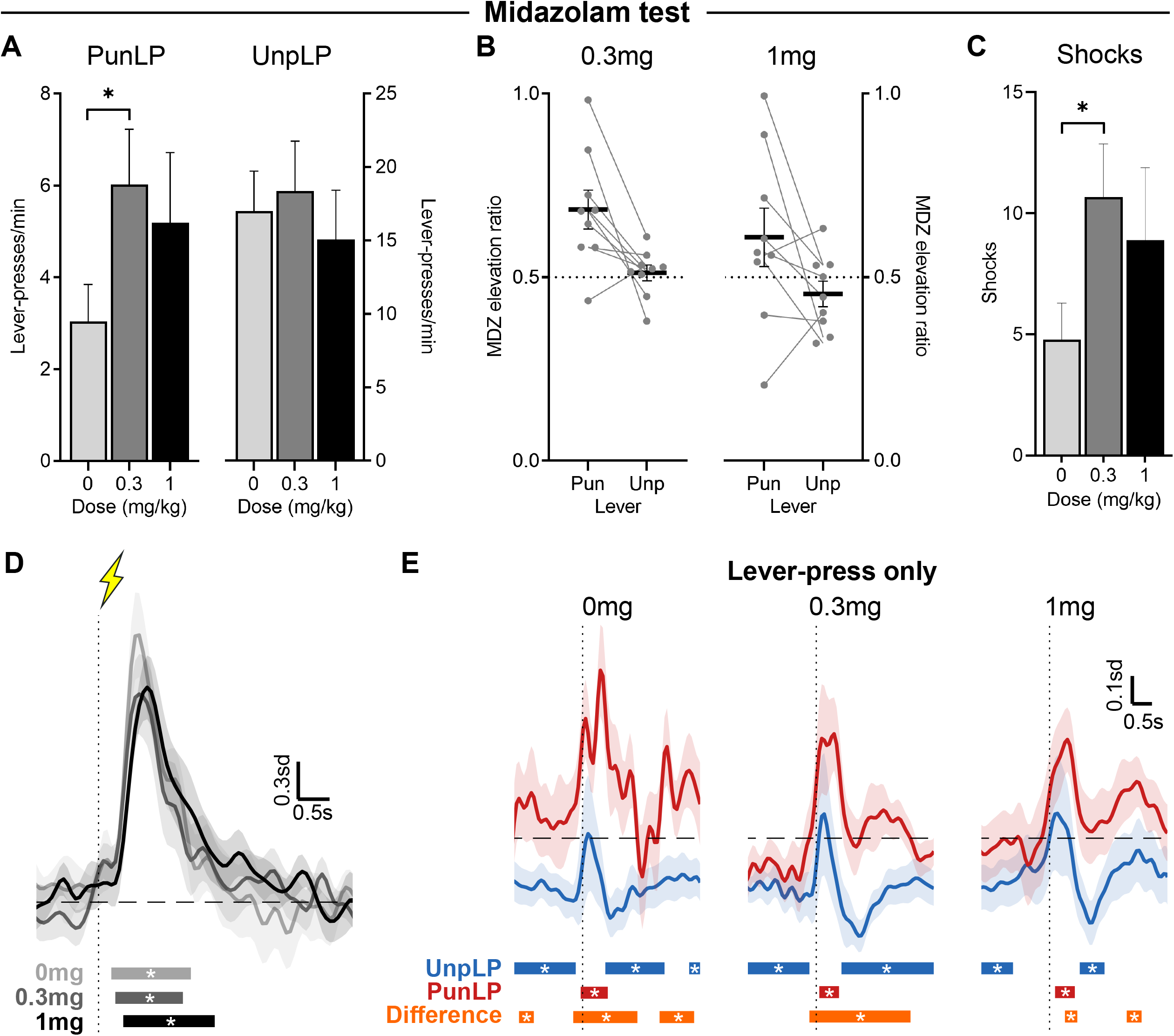
Influence of midazolam (MDZ) on punished behavior and BLA activity. **[A]** Mean±SEM punished (*left*) and unpunished (*right*) lever-press rate across MDZ doses (*n* = 9). **[B]** Elevation ratio of lever-pressing (relative to 0mg/kg) for 0.3mg/kg (*left*) and 1mg/kg (*right*) MDZ; bar = mean±SEM, dots = individuals. **[C]** Mean±SEM shocks received across MDZ doses. **[D]** Mean±SEM BLA activity kernel around shock deliveries across MDZ doses. Bars at bottom of graph indicate significant deviations from baseline (dF/F ≠ 0). Vertical dashed line indicates shock onset, horizontal dashed line indicates baseline (dF/F = 0). **[E]** Mean±SEM activity kernel around punished (PunLP; red) and unpunished (UnpLP; blue) lever-presses (no outcome) across MDZ tests. Bars at bottom of graph indicate significant deviations from baseline (95% CI) for PunLP and UnpLP, and significant differences between punished and unpunished kernels (orange). Vertical dashed lines indicate time of lever-press, horizontal dashed lines indicate baseline (dF/F = 0).

Although there was a trend towards increased punished lever-press rates for both doses of MDZ relative to control (Figure 3A), only 0.3mg caused a significant and selective increase in punished responding (F(1,8) = 19.93, *p* = .002); 1mg did not robustly increase punished responding (F(1,8) = 1.12, *p* = .320). There was no significant effect of 0.3mg (F(1,8) = 0.727, *p* = .419) or 1mg (F(1,8) = 1.738, *p* = .224) MDZ on unpunished lever-press rates.

The effect of MDZ on lever-pressing was also assessed via an elevation ratio (Figure 3B). This normalizes MDZ lever-press rates against control lever-press rates, providing a more sensitive measure of directional, proportional change. When comparing ratios against the null of 0.5 (no difference), 0.3mg MDZ significantly increased punished lever-pressing (t(8) = 3.485, p = .008), without commensurately affecting unpunished lever-pressing (t(8) = 0.5282, p = .612). As found using lever-press rates, 1mg MDZ did not reliably increase punished lever-pressing (t(8) = 1.365, p = .209) or decrease unpunished lever-pressing (t(8) = −1.327, p = .221). When considering the substantial individual differences in effects of 1mg MDZ, the two subjects that showed decreased punished lever-pressing at 1mg MDZ also showed decreased unpunished pressing. This pattern is consistent with general response reduction produced by sedating doses of MDZ (Pieri, 1983). These two subjects showed increased punished lever-pressing at 0.3mg MDZ, but may have been particularly sensitive to MDZ, causing sedative effects to dominate at 1mg/kg.

These impacts of MDZ on punished behavior were consequential. The increased punished lever-pressing following 0.3mg MDZ resulted in significantly more shocks delivered (F(1,8) = 19.99, *p* = .002).

Interestingly, shock evoked BLA transients (Figure 3D) were unaffected by MDZ, suggesting that the anti-punishment effects of MDZ were not due to a change in BLA encoding of the punisher per se. Indeed, across MDZ tests, shocks elicited robust excitatory transients of similar magnitude and duration to those observed in non-MDZ punishment sessions. Punished and unpunished lever-pressing were associated with excitatory and inhibitory BLA transients, respectively. However, excitation around the punished lever-press, particularly that immediately preceding the lever-press, was less pronounced under MDZ.

#### Assessing changes in BLA activity across sessions

To directly compare peri-event BLA activity across punishment learning and MDZ tests, we calculated the fit between punished lever-press, unpunished lever-press, and shock waveforms across sessions. Fit scores quantify the degree to which one waveform is similar to another: positive values indicate matching transients (1 indicates identical waveforms relative to baseline) whereas negative values indicate opposite transients (−1 indicates perfectly inverse waveforms relative to baseline). Significant positive and negative fits were identified through fit confidence intervals bootstrapped from subject mean waveforms. Fits between the various waveforms are depicted in Figure 4A. The similarity/dissimilarity between waveforms was also depicted in 2D space (Figure 4B), such that similar waveforms are plotted closer together than dissimilar waveforms. Grey lines connecting datapoints in Figure 4B indicate significantly positive fits, i.e. activity pattern clusters.

**Figure 4.**
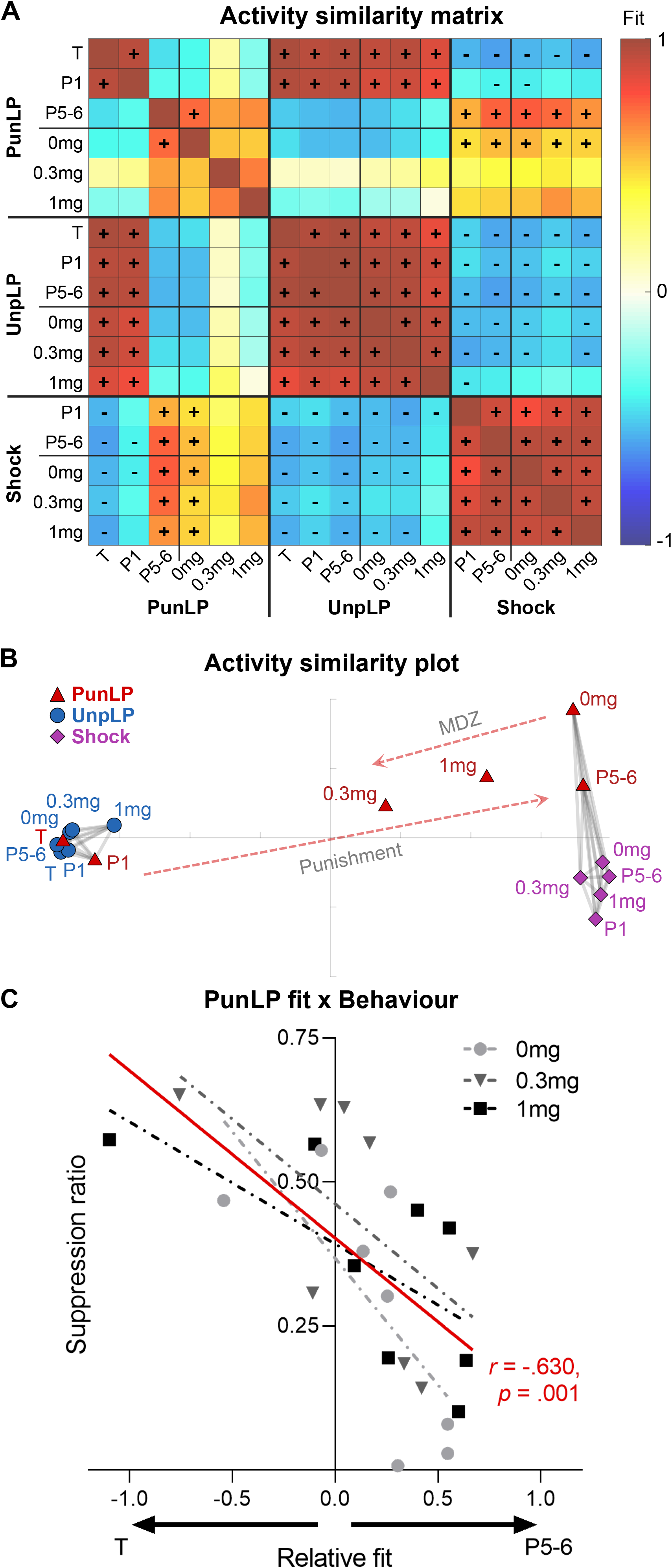
Change in BLA activity across punishment and midazolam tests. **[A]** Similarity of punished lever-press (PunLP), unpunished lever-press (UnpLP) and shock activity kernels across lever training (T), punishment (P1, P5-6), and MDZ tests (0mg, 0.3mg, 1mg), determined via normalized fit between mean of subject waveforms. Significant positive (+) and negative (-) fits were identified via bootstrapping (95% CI of fits). UnpLP and shock-related activity kernels are relatively unchanged throughout punishment and MDZ tests, whereas PunLP kernels change substantially. **[B]** 2D plot of kernel similarities (multidimensional scaling of fit distance [1 – fit]). Gray lines connect significantly similar waveforms. Dashed arrows indicate the general effect of punishment and MDZ on PunLP coding in BLA. Punishment transformed PunLP kernels from UnpLP-like kernels into shock-like kernels; MDZ reverted PunLP kernels towards its pre-punishment state. **[C]** Relationship between subject’s BLA PunLP activity and PunLP behavior across MDZ tests (0mg, 0.3mg, 1mg). Fit of MDZ test PunLP activity onto late punishment vs. pre-punishment PunLP activity was quantified in a relative fit score (P5-6 fit – T fit, per subject). Relative fit was compared against PunLP suppression (suppression ratio relative to T, per subject): less punishment avoidance was associated with PunLP activity kernels being more similar to pre-punishment than late punishment.

Unpunished lever-presses and shock transient waveforms were stable across punishment and MDZ test sessions. Cross-session fits of unpunished lever-press transients were uniformly high and significantly similar (Figure 4A; mean fit = 0.938), as were cross-session fits of peri-shock transients (mean fit = 0.914). When comparing fits between unpunished lever-press and shock transients, fits were reliably negative (mean fit = −.547), reflecting the inhibitory versus excitatory transients observed around these respective events across sessions. Visualizing these relationships in a 2D similarity plot (Figure 4B) showed that unpunished lever-press transients across sessions occupied a tightly clustered space, while shock transients occupied a separate but similarly tightly clustered space.

In contrast, punished lever-press transients changed substantially across punishment and MDZ sessions. During training and first punishment session, transients around punished lever-presses were highly similar to unpunished lever-pressing and dissimilar to shock transients. However, for later punishment sessions, the reverse became true; punished lever-press transients diverged from unpunished lever-press transients and became significantly similar to shock transients, as highlighted by the rightward arrow in Figure 4B.

Importantly, MDZ reverted punished lever-press transients towards their pre-punishment state. Punished lever-press transients following control (0 mg) injections were similar to late punishment lever-press and cross-session shock transients. This similarity was undermined by MDZ, with punished lever-press activity following 0.3 mg and 1 mg MDZ no longer showing significant similarity to those transients (Figure 4A,B). Critically, punished lever-press transients became more similar to pre-punishment/unpunished lever-press transients under MDZ, as highlighted by the leftward arrow in Figure 4B. This was particularly true for 0.3 mg punished lever-press transients, whose fit against unpunished and pre-punishment lever-press activity trended positive instead of negative (Figure 4A). It is noteworthy that this stronger effect for 0.3 mg MDZ on BLA punished lever-press activity mirrors the stronger effect of this dose on punished lever-pressing itself.

#### Relationship between anti-punishment effects of MDZ and BLA lever-press transients

These analyses indicate that MDZ increased punished responding and reverted BLA coding of punished behavior towards a pre-punishment state. So, we investigated the direct relationship between these two effects, particularly given that the anti-punishment effects of MDZ on behavior appeared to be influenced by individual differences.

To address this, we first calculated a relative fit score that quantified how strongly each subject’s punished lever-press transients per MDZ test (0mg, 0.3mg, 1mg) fit their punished lever-press activity during late punishment versus pre-punishment. We found a strong negative relationship (*r* = −.63, *p* = .001) between how much punished lever-press transients reverted to their pre-punishment form and how much punished behavior reverted to pre-punishment levels. In other words, the more that punished lever-press transients reverted to their pre-punishment form, the more animals failed to avoid punishment. This inverse relationship was true across MDZ doses, suggesting the anti-punishment effects of MDZ are mediated by, or at least tracked by, the degree to which MDZ influences BLA coding of punished responses.

## Discussion

Punishment involves learning the instrumental contingencies between actions and their adverse consequences. Here we used a well-controlled, within-subjects punishment task to study how BLA supports punishment learning. Rats were trained to respond on two levers for food reward prior to one of those responses being punished with footshock. Punishment was effective. It caused response-specific suppression of lever-pressing, increased the latencies with which rats responded on the punished lever, but did not increase levels of immobility indicative of involuntary conditioned fear responses.

Using fiber photometry, we examined BLA principal neuron Ca^2+^ transients during instrumental punishment and found that BLA activity encodes instrumental aversion. At the population-level, BLA principal neurons exhibited phasic excitatory transients to response-elicited footshock punishers across punishment sessions. Prior to punishment BLA principal neurons exhibited phasic inhibitions to lever-presses associated with reward. However, across punishment sessions, BLA transients to the punished action became excitatory whereas unpunished actions remained inhibitory. This same profile of within-subject, bidirectional BLA transients was observed during choice tests when the punisher itself was absent. So, there was evidence here for BLA encoding of both instrumental aversive outcomes and specific, punished instrumental actions. This specific BLA encoding of both punished instrumental actions and their consequences supports the view that BLA is a critical neural substrate for instrumental aversion (Jean-Richard-Dit-Bressel et al., 2018; Killcross et al., 1997b; Piantadosi et al., 2021).

Our findings suggest that punished responses may evoke punisher-specific representations in BLA to guide behavior. Similarity analyses showed that BLA Ca^2+^ transients to punished instrumental actions became highly similar to the punisher itself across training. This parallels findings from Pavlovian fear conditioning where BLA ensemble activity to Pavlovian CSs becomes more similar to the shock US outcomes they predict (Grewe et al., 2017; Zhang & Li, 2018). In Pavlovian tasks, increased alignment of BLA activity between antecedents and outcomes has been interpreted in two distinct ways. First, it may reflect common behavioral responses to CSs (i.e. antecedents) and USs (i.e. outcomes) as they come to demand increasingly similar behavioral outputs across conditioning (Kyriazi et al., 2018). According to this view, the similarity in BLA Ca^2+^ transients to punished lever-presses and footshock reflect underlying similarity in behavioral responses to them. Although this seems plausible in Pavlovian tasks, it is an unlikely explanation of the present data because lever-presses are not responses to footshock. Second, this increased alignment could reflect BLA encoding of the aversive value common to punished actions and punishers. Indeed, consistent with this, BLA Ca^2+^ transients throughout the task were well-represented in a low dimensional space corresponding to a continuum of aversive valence (Figure 4). Representations of aversive value are presumed necessary for appropriate punishment avoidance, and interfering with BLA activity, including during the moments of punishment, reduces avoidance (Jean-Richard-Dit-Bressel & McNally, 2015; Killcross et al., 1997b; Orsini et al., 2017; Orsini et al., 2015; Pelloux et al., 2013; Piantadosi et al., 2017; Verharen et al., 2019). Given that punishment learning is putatively underpinned by specific Action–Punisher associations (Bolles et al., 1980; Jean-Richard-Dit-Bressel et al., 2018), our findings suggest that punished responses may evoke punisher-specific representations in BLA to guide behavior, a function already ascribed to BLA for appetitive outcomes (Balleine et al., 2003).

The results from midazolam tests strengthen this interpretation. Systemic administrations of midazolam not only increased punished responding but they also reverted BLA activity around punished responses towards their unpunished state. Critically, BLA activity around footshock and unpunished responses were relatively unaffected by midazolam. So, the selective effects of midazolam on punishment were linked to selective effects on BLA coding of punished actions. This shows that benzodiazepines do not undermine aversion coding in BLA generally (Escorihuela et al., 1993; McNaughton & Gray, 2000), but rather they interfere with the response-specific representations promoting punishment. Critically, individual differences in the effects of midazolam on punishment avoidance were directly predicted by the degree to which the benzodiazepine could revert lever-press activity to its pre-punishment state. This suggests that individual differences in the anxiolytic action of benzodiazepines, traditionally assessed using punishment tasks (Pollard & Howard, 1990), may be linked to individual variation in this reversion.

There are three methodological limitations worth considering. First, the population readout obtained here using fiber photometry prevents inferences about the activity of individual BLA neurons. BLA neurons can exhibit marked activity heterogeneity to stimuli and behaviors (Carelli et al., 2003; Kyriazi et al., 2018). Moreover, both the activity and functions of BLA principal neurons can be dissociated according to their specific projection targets (Janak & Tye, 2015). So, whether and how the excitatory and inhibitory activity we observed to outcomes and punished versus unpunished responses were driven by overlapping or separate neuronal ensembles remains unclear. How punishment is encoded across BLA ensembles, and how these in turn influence broader network activity to guide behavior, is an important area for future research. Second, we administered midazolam systemically, so the effects observed here may be mediated by actions in brain regions beyond BLA. It is worth emphasizing that the effect of benzodiazepines on punishment avoidance are linked to, and directly recapitulated by, its actions within BLA (Hodges et al., 1987; Liu & Glowa, 2000). However, other sites of action such as hippocampus may contribute to the effects of benzodiazepines on behavior and BLA activity (File, 2000; McNaughton & Gray, 2000; Tan et al., 2010). Third, we studied only male rats here, so the nature, role, and relevance of sex differences in punishment remains worth investigating. It is worth noting sex differences using immediate punishment tend to be modest and attributable to increased propensity to fear over punishment learning in females (Chowdhury et al., 2019; Jean-Richard-Dit-Bressel et al., 2019; Liley et al., 2019; Orsini et al., 2016). Moreover, our rodent work, based largely on male rats, accurately accounts for punishment learning in female and male humans (Jean-Richard-Dit-Bressel et al., 2021).

In summary, we investigated BLA principal neuron activity across punishment learning, choice, and under the influence of benzodiazepine. We show that instrumental punishment is encoded in BLA activity via excitations to punishers and their behavioral antecedents, in contrast to inhibitions around unpunished actions. Benzodiazepine increased punished behavior, but only to the extent it reverted BLA coding of punished actions to their pre-punishment state. Together, these findings show that punishment learning is supported by aversion-coding of instrumental responses in the BLA and that the anti-punishment effects of benzodiazepines are associated with a reversion of this aversion coding.

## Author contributions

<u>Conceptualisation</u>: PJRDB, GPM. <u>Programming and Experimentation</u>: PJRDB, JT, AD. <u>Formal Analysis</u>: PJRDB, JT, GPM. <u>Writing – original draft:</u> PRDB, GPM. <u>Writing – Review & Editing</u>: All authors. <u>Funding acquisition</u>: GPM.

